# Aberrant TNF signaling in pancreatic lymph nodes of patients with Type 1 Diabetes

**DOI:** 10.1101/2024.05.31.596885

**Authors:** Maryam Abedi, Priyadarshini Rai, Yeqiao Zhou, Chengyang Liu, Isabelle Johnson, Aditi Chandra, Maria Fasolino, Susan Rostami, Klaus H. Kaestner, Ali Naji, Robert B. Faryabi, Golnaz Vahedi

## Abstract

The therapeutic landscape for Type 1 Diabetes (T1D) is rapidly changing as ongoing clinical trials aim to delay beta-cell loss by inhibiting proinflammatory cytokines. However, the precise timing and cellular contexts of cytokine dysregulation remains unknown. We generated the largest existing measurement of gene expression and chromatin accessibility in ∼1 million immune cells from the pancreatic lymph nodes and spleens of 34 T1D and non-diabetic organ donors. Our study revealed heightened gene activity of the tumor necrosis factor (TNF) pathway and subsequent chromatin remodeling in central memory CD4^+^ T cells residing in the pancreatic lymph nodes of T1D and non-diabetic islet-autoantibody positive donors. These findings, validated in mice, offer a mechanism underlying the efficacy of TNF inhibitors, currently undergoing clinical trials to delay T1D onset.

## Introduction

Insulin, which is a hormone produced by pancreatic islet beta cells, facilitates glucose uptake by peripheral tissues such as adipose tissues and skeletal muscles. In the autoimmune disease Type 1 Diabetes (T1D), T lymphocytes assault and destroy beta cells. Genetics plays a strong role in determining T1D risk although ill-defined environmental factors trigger disease onset. Genome-wide association studies (GWAS) have pinpointed genomic regions containing single-nucleotide polymorphisms (SNPs) associated with T1D risk^1,2^. Variations in major histocompatibility complex (MHC) class II genes, which are responsible for presenting peptides to CD4^+^ T cells, contribute to approximately half of the genetic predisposition to T1D^2^. Apart from the MHC region, more than 100 SNPs mostly within the noncoding genome have been reported to increase the risk for T1D^1,2^. Despite these findings, the precise origin of this autoimmune condition remains unknown.

Insulin replacement has stood as the primary therapy for T1D patients for more than a century^3^. Nonetheless, the therapeutic landscape to delay or prevent T1D is rapidly changing as multiple ongoing clinical trials are examining immunotherapeutic approaches targeting cytokines, chemokines, or selectively controlling specific immune cell types. The first immune modulatory drug which received Food and Drug Administration (FDA) approval in 2022 is teplizumab^4,5^, an anti-CD3 monoclonal antibody affecting T cell function. Teplizumab can delay the T1D onset in individuals with preclinical characteristics by two years^4^ and has clinical benefits in recently diagnosed T1D patients^5^. Clinical trials targeting the tumor necrosis factor alpha (TNF-a) pathway using golimumab^6,7^ or etanercept^8^ demonstrated improved clinical indicators in recently diagnosed patients. An initial trial of baricitinib, which blocks Janus Kinase 1 (JAK1) and JAK2 required for cytokines signal transduction, also showed improved beta-cell function in newly diagnosed individuals^9^.

Although the inhibition of proinflammatory cytokines holds the promise for favorable clinical outcomes in T1D, there remains a profound lack of understanding concerning the precise timing and specific cellular contexts in which dysregulated cytokine expression contributes to the progression of autoimmunity. Additionally, the lack of reliable biomarkers at early stages of T1D makes it difficult to pinpoint pre-symptomatic individuals who could benefit from advanced immunomodulation techniques. The current standard for early detection of autoimmunity relies on identifying and quantifying the number, type, and titer of specific islet autoantibodies present in the serum of at-risk individuals^10^. However, islet autoantibodies alone are not robust early predictors of individual progression to T1D onset^11^. Deep profiling efforts that can delineate the precise timing and specific cellular contexts in which dysregulated cytokine expression contributes to the progression of autoimmunity hold the potential to profoundly reshape the therapeutic outlook for T1D.

We reasoned that molecular profiling of immune cells in pancreatic related tissues could unravel the identity of immune subsets receiving specific cytokine signals during the disease progression. This goal became feasible thanks to the NIDDK-supported Human Pancreas Analysis Program (HPAP)^12,13^, which is an ongoing effort to collect pancreas and immune-related tissues from hundreds of organ donors. As part of this program, we received pancreatic lymph nodes (PLNs) from 13 non-diabetic controls, 8 non-diabetic but islet autoantibody positive (AAb^+^), and 13 T1D organ donors. We focused our deep immune profiling on cells residing in PLN, a critical site for the drainage of immune cells into the pancreas and priming of autoreactive T cells^14,15^. We also collected the spleen tissues from the same individuals with the goal of narrowing down immunological perturbations unique to the pancreatic tissues. Since the chromatin structure is an attractive template for transmitting and remembering cytokine signals^16–18^, we chose the multiome single-cell profiling assay^19^ which allowed us to measure chromatin accessibility and gene expression in the same cell across more than 1 million cells collected from PLNs and spleens of 34 organ donors.

In this work, we report strong activation of tumor necrosis factor (TNF) cytokine families inducing the expression of NF-kb transcription factor family and chromatin remodeling associated with this transcription factor in central memory CD4^+^ T cells residing in PLNs of T1D and AAb^+^ organ donors. During T1D pathogenesis, islet protein peptides loaded onto MHC class II proteins in dendritic cells engage central memory CD4^+^ T cells^20^, which have the unique ability to replenish immune responses upon antigen encounters^21^. This reactivation of memory CD4^+^ T cells produces the cytokines required to activate or ‘license’ CD8^+^ cytotoxic T cells, leading to beta cell killing. As discovered here, the activation of central memory CD4^+^ T cells occurs in the PLNs of T1D patients as well as normoglycemic individuals who present with circulating autoantibodies. Remarkably, single-cell profiling in the animal model of T1D validated these findings. The unexpected activation of TNF pathway in AAb^+^ donors provides a plausible mechanism of action for golimumab, a TNF neutralizing monoclonal antibody which is being evaluated to delay disease onset in phase II clinical trials^6,7^. Golimumab may prevent central memory CD4^+^ T cells in PLNs from helping CD8^+^ T cells, delaying beta cell loss in pre-symptomatic individuals.

## Results

### Immune cell composition in pancreatic lymph nodes

In this study, we defined the molecular signatures of immune cells in pancreatic lymph nodes (PLNs), which are thought to harbor the cells engaged in the initial autoimmune assault on beta cells^14,15,22^. We obtained PLNs collected by the HPAP team from 13 individuals clinically diagnosed with T1D, 8 non-diabetic individuals positive for islet autoantibodies (AAb^+^), and 13 individuals as healthy controls (HD) (Figure 1A-B). Clinical information related to donors used in our study is provided in Figure 1B and Table S1. Additional clinical details collected by HPAP investigators along with the molecular data generated in this study are shared on the open-access database called <u>PANC-DB</u>. When feasible, our surgical team collected PLN tissues from different anatomical sections, i.e. the head, body, and tail of the pancreas. Considering the variability in the quality of pancreatic tissues after surgery, the complete set of head, body, and tail regions is available in only a subset of HPAP donors. Tissues from the head of the pancreas were available for most donors included in our study (Figure 1B). In our analytical strategies to define differentially expressed genes or differentially accessible regions, we either combined cells from different PLN compartments or used cells from the pancreas’ head separately.

**Figure 1.**
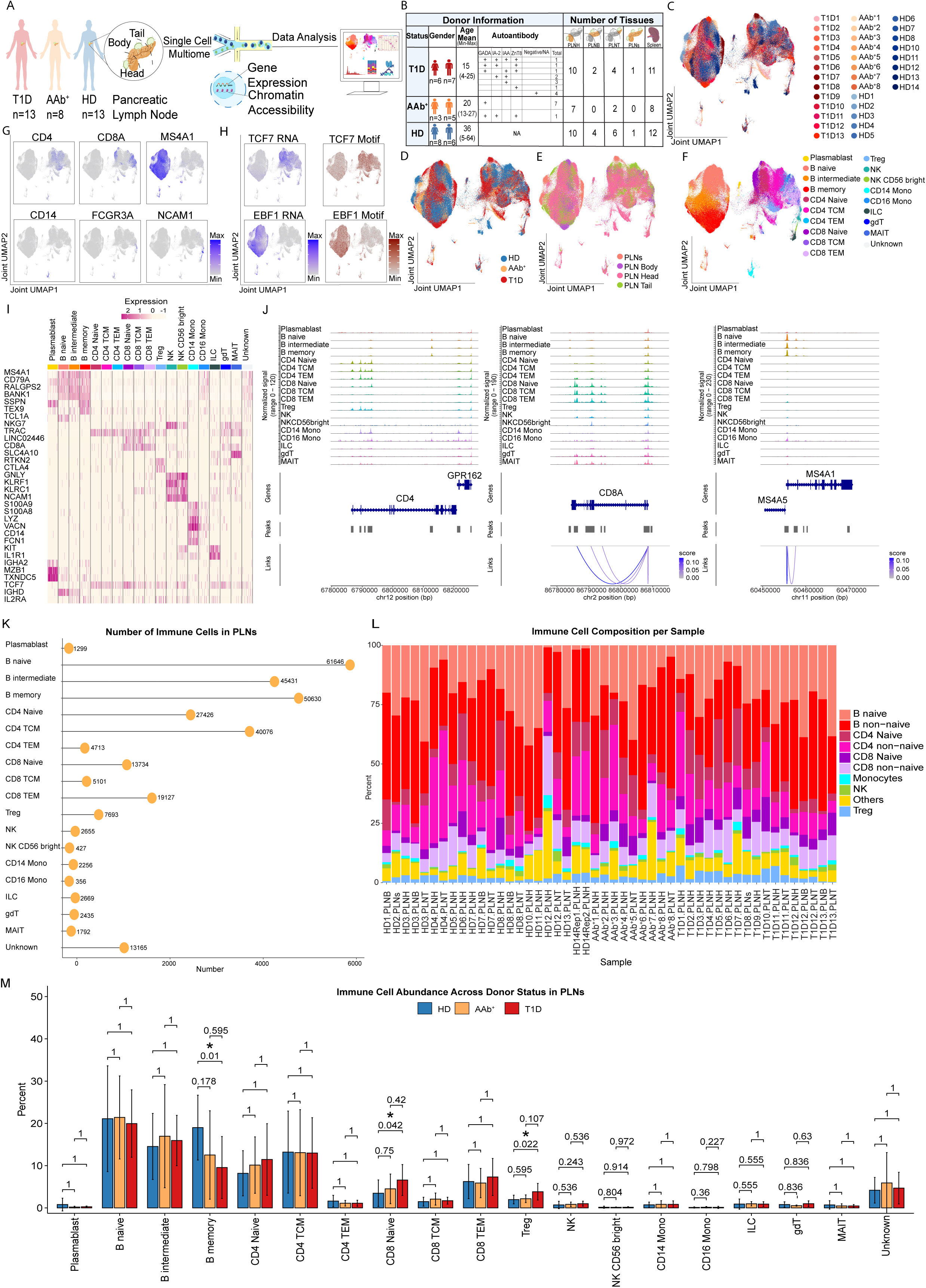
The immune cell atlas of PLNs in T1D, AAb^+^, and HD organ donors. (**A**) Schematic outlining the experimental approach in this study for molecular profiling of immune cells in the PLN tissues. (**B**) The clinical and demographic information of donors in this study. (**C-F**) The Uniform Manifold Approximation and Projection (UMAP) representing immune cells remained after quality control analysis labeled with donor number as listed in Table S1 (**C**), donor status (**D**), tissue types (**E**), and immune cell annotation (**F**). (**G**) UMAPs representing RNA levels of cell-type specific markers. (**H**) UMAPs representing RNA levels of B and T cell-specific transcription factors in addition to motif analysis in cCREs. (**I**) The heatmap showing the gene expression of cell markers employed for immune cell annotation. (**J**) The coverage plot indicating chromatin accessibility in gene marker location corresponding to the cell annotation. (**K**) The lollipop plot showing the number of cells per annotated immune cell type after quality control analysis. (**L**) The stacked bar chart indicating the immune cell composition in each donor. (**M**) The bar chart showing immune cell abundance for cells grouped in different donor groups. Asterisks represent adjusted p-value ≤ 0.05. T1D: Type 1 diabetes, AAb^+^: autoantibody positive, HD: healthy control donor, GADA: glutamic acid decarboxylase antibody, IA-2: Islet antigen 2, IAA-2: insulin autoantibody, ZnT-8: zinc transporter 8. In (**B**) PLNH: pancreatic lymph node from the pancreas head, PLNB: pancreatic lymph node from the pancreas body, PLNT: pancreatic lymph node from the pancreas tail.

We sought after a molecular profiling assay which was capable of simultaneously measuring chromatin accessibility and gene expression in the same cell across thousands of individual cells per donor. We reasoned that it is essential to annotate cells using the gene expression modality and define epigenomic scars caused by the cytokine milieu using the chromatin accessibility modality. Hence, we conducted multimodal single-nucleus (sn)ATAC/snRNA-seq experiments in 565,887 cells obtained from 46+ PLN samples in 34 HPAP donors (Figure S1A-C). The two modalities were integrated using weighted nearest neighbor (WNN) analysis^19^. Subsequently, the uniform manifold approximation and projection (UMAP) was employed on the WNN graph and clustering was performed (Figure S1A-E). We removed any potential doublets (Figure S1F) and further filtered cells which did not pass the quality control for both snRNA and snATAC assays (Figure S1G-I). After this stringent quality control, we retained 302,631 individual cells with high quality in both RNA and ATAC modalities and observed mixing of cells across donors, disease status, and anatomical PLN regions (Figures 1C-F and S1J-K). This outcome underscores the robustness and high quality of our multimodal dataset. For cell type annotation, we followed a unified strategy for reference assembly and transfer learning based on the RNA modality, reporting 30 distinct cell types in the innate and adaptive immune compartments^23^ (Table S2). We further labeled 4.35% of cells as “Unknown” considering the low confidence of our cell annotation strategy in this small subset (Figures 1F and S1E). Visualizing the expression of marker genes across different annotated cell types on UMAP or heatmap in addition to coverage plots of chromatin accessibility at cell-type-specific genomic loci corroborated the high quality of our annotation strategy (Figures 1G-J and S1L). For example, both mRNA abundance and chromatin accessibly of the *MS4A1* gene, encoding CD20 a B-lymphocyte-specific membrane protein, were confined to B cells. The unknown cells and annotated cell types with less than 100 cells were excluded from further analysis which left 18 major cell types in the innate and adaptive immune compartments in PLN tissues (Figures 1K and S1J-K). In line with prior reports^24^, B and T lymphocytes were the most frequent cell types while CD16^+^ monocytes were less abundant in PLNs (Figure 1K). We next identified similarities and differences for cell composition across individual donors (Figures 1L) and also compared cell type frequencies for cells grouped as T1D, AAb^+^, and HD (Figures 1M and S1M). Our investigation did not yield substantial evidence indicating significant alterations in global cell composition associated with T1D. Yet, we found moderate differences in the frequency of some cell types such as naïve CD8^+^ T cells, CD4^+^ T regulatory cells (Tregs), naive CD4^+^ T cells, and memory B cells in PLNs of T1D donors compared with other donor groups (Figures 1M and S1M). Together, we generated the largest existing collection of multi-modal molecular data for human PLNs.

### Immune cell composition in spleens

To delineate the PLN-specific immune signature associated with T1D, we also measured the molecular landscape of CD45^+^ immune cells in the spleen, which is a secondary lymphoid organ situated distally from the pancreas. We conducted snATAC/snRNA-seq multiome experiments in 459,983 cells on samples collected from the spleens of the same donors that PLNs were collected (Figures 2A and S2A-E). We followed a similar analytical strategy as described above to process samples collected in spleens. After we applied stringent quality control, we retained 328,855 individual cells with high-quality RNA and ATAC modalities (Figure S2F-I). Dimensionality reduction with UMAP using cells from spleens demonstrated mixing of cells across donors and disease status, suggesting the high-quality of our multimodal data (Figure 2B-D). We followed a similar cell annotation strategy as described for PLNs and defined different immune populations (Figures 2D and S2J-K). The expression of marker genes and coverage plots of chromatin accessibility at cell-type-specific genomic loci also confirmed the high-quality of our cell annotation strategy in spleens (Figures 2E-H and S2L). We labeled 1.19% of cells as “Unknown” considering the low confidence of our cell annotation strategy in this small subset and excluded these cells along with the cell types with less than 100 cells from further analysis (Figure S2E). Because of the larger number of cells available from spleens as opposed to PLNs, we were also able to perform the trimodal assay termed ‘TEA-seq’^25^ (Transcription, Epitopes, and Accessibility) on a subset of our spleen samples. In this assay, determination of transcript abundance and chromatin accessibility is combined with limited proteomics to further refine cell type annotation^25^. Integration of cells from a subset of samples with protein expression measured by the trimodal assay confirmed our overall strategy for cell type annotation based on RNA expression and transfer learning (Figure 2I-K).

**Figure 2.**
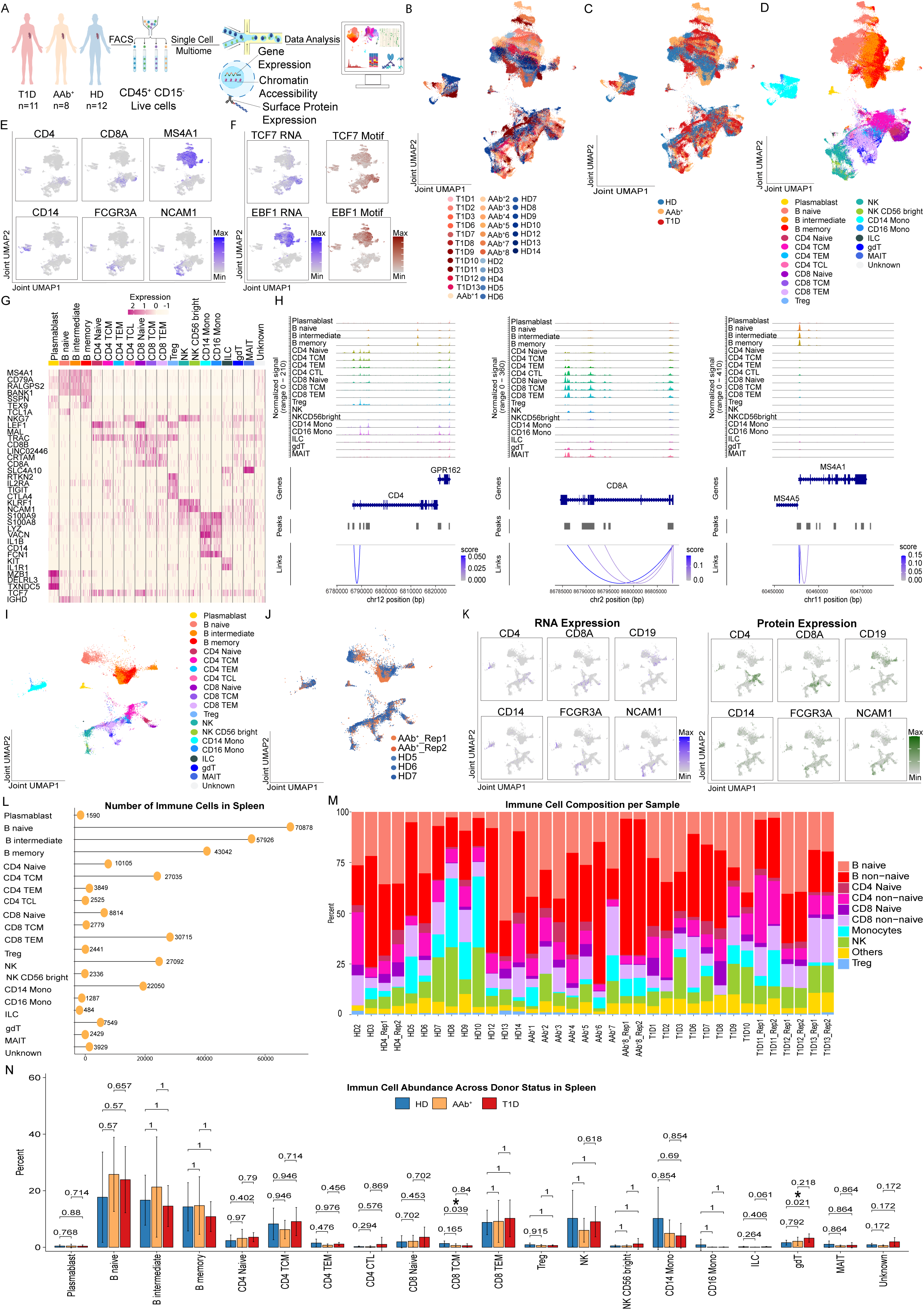
Immune cell atlas of spleens in T1D, AAb^+^, and HD organ donors. (**A**) Schematic outlining the experimental approach in this study for molecular profiling of immune cells in the spleen tissues. (**B-D**) UMAP representing immune cells remained after quality control analysis labeled with donor number as listed in Table S1 (**B**), donor status (**C**), and immune cell annotation (**D**). (**E**) UMAPs representing RNA levels of cell-type specific markers. (**F**) UMAPs represent RNA levels of B and T cell-specific transcription factors in addition to motif analysis in cCREs. (**G**) The heatmap showing the gene expression of cell markers employed for immune cell annotation. (**H**) The coverage plot indicating chromatin accessibility in the gene marker location corresponding to the cell annotation. (**I-J**) The UMAP representing a subset of samples where TEA-seq have been generated for them containing protein modality labeled with immune cell annotation (**I**) and donor number (**J**). (**K**) The feature plots demonstrating the gene expression of cell markers and protein expression of the same markers in the subset of samples. (**L**) The lollipop plot showing the number of cells per annotated immune cell type after quality control analysis. (**M**) The stacked bar chart indicating the immune cell composition in each donor. (**N**) The bar chart showing immune cell abundance between donor status. Asterisks represent adjusted p-value ≤ 0.05. T1D: Type 1 diabetes, AAb^+^: Autoantibody positive, HD: Healthy control donor.

Overall, spleen samples were composed of B and T lymphocytes, NK cells, and CD14^+^ monocytes, alongside a spectrum of other immune subpopulations detected at varying frequencies (Figure 2L). Although cell composition across samples demonstrated similarities and differences across individual donors (Figure 2M), only moderate differences in cell composition of some cell types such as gamma-delta T cells and central memory CD8^+^ T cells were detected in spleen tissues of T1D donors (Figures 2N and S2M). Together, the collection of multi-dimensional molecular data in spleens and PLNs of human organ donors provided us with the opportunity to define PLN-specific molecular signatures of T1D progression.

### Large-scale changes in transcriptional outputs of central memory CD4^+^ T cells in T1D PLNs

To investigate whether the RNA modality can detect genes whose expression are deregulated across T1D donors, we performed pseudo-bulk differential gene expression analysis between donor groups for the major annotated cell types separately for PLNs and spleens. Using stringent criteria (|log2 foldchange| > 0.5 and adjusted p-value < 0.05), we identified the number of genes whose expression increased or decreased in pair-wise comparisons between T1D vs HD, T1D vs AAb^+^, and AAb^+^ vs HD donors. In both PLN and spleen tissues, the comparison of cells between T1D and HD donors demonstrated the largest difference in gene expression levels (Figure 3A-B, red and dark-blue bars). Although cells from AAb^+^ and HD donors were also transcriptionally different, the T1D vs AAb^+^ comparison demonstrated minimal number of differentially expressed genes for most cell types, implying large-scale similarity in transcriptional landscapes of immune cells in PLNs of T1D and AAb^+^ donors (Figure 3A-B). Remarkably, the comparison of various cell types in PLN samples highlighted central memory CD4^+^ T cells to be the most transcriptionally distinct cell type between T1D and HD donors (Figure 3A). The heatmap visualization of gene expression levels from central memory CD4^+^ T cells by donor type corroborated their differential expression across donors (Figure 3C). Although most of these genes did not pass the statistical threshold to be referred to as differentially expressed between AAb^+^ and HD donors, the heatmap visualization of the expression levels of these genes suggested the similarity of AAb^+^ to T1D donors in comparison to HD counterparts (Figure 3C). In the spleen, memory B cells displayed the largest change in gene expression levels in the T1D vs HD comparison (Figure 3B). The comparison of genes across the three pair-wise comparisons suggested the uniqueness of genes in the T1D vs HD comparison across different tissues (Figure S3 A-C). Together, our side-by-side comparison of cells obtained from different tissues emphasized the importance of analyzing PLNs for a thorough understanding of T1D progression.

**Figure 3.**
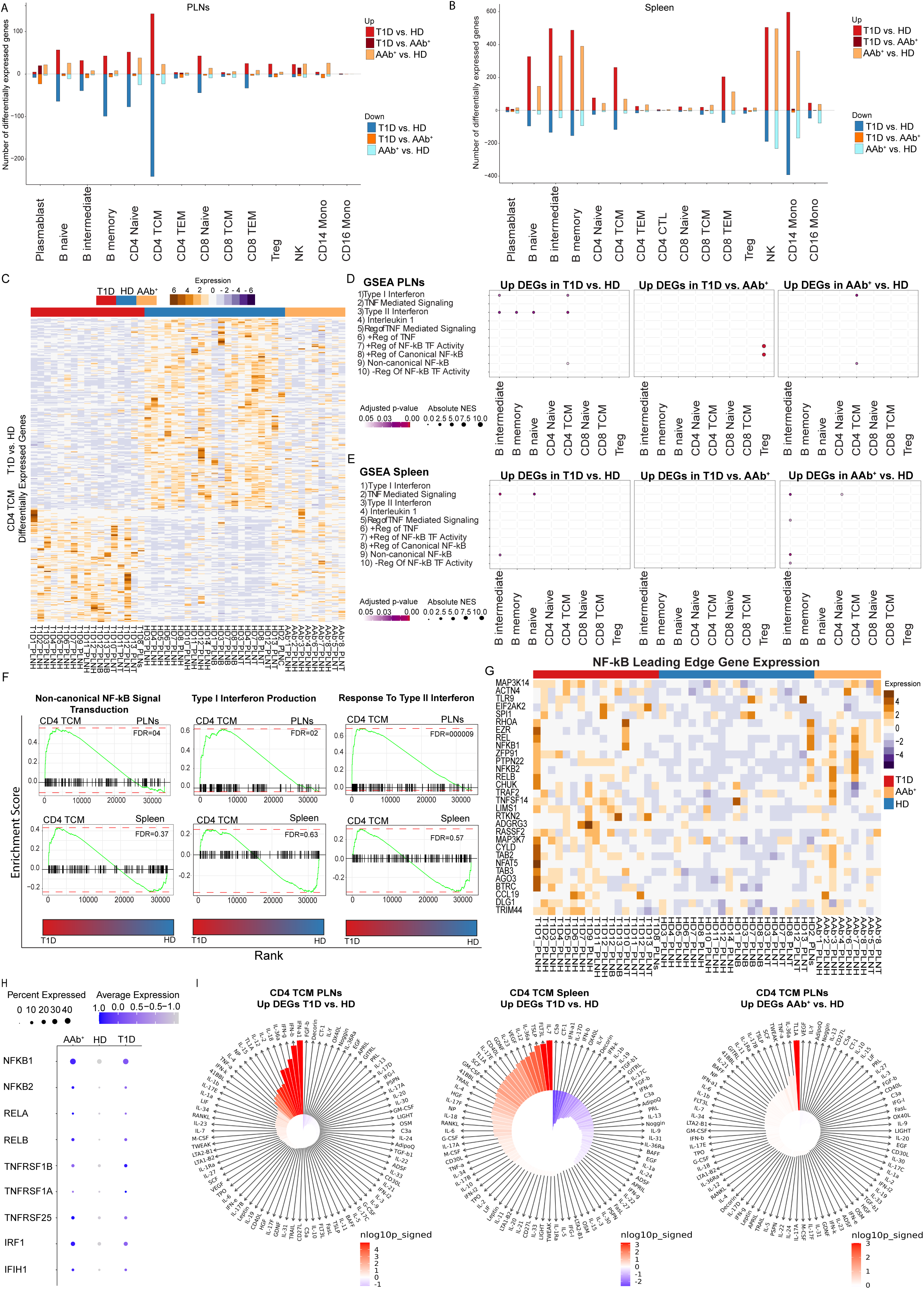
TNF and interferon signaling are transcriptionally dysregulated in central memory CD4^+^ T cells in PLNs. (**A-B**) The bar plot demonstrating the number of differentially expressed genes in each cell type between T1D vs HD, T1D vs AAb^+,^ and AAb^+^ vs HD in PLNs (**A**) and spleen (**B**). For differential analysis, first pseudobulk data was created based on the cell type and donor ID. Subsequently, *edgeR LRT* (edgeR likelihood ratio test)^44^ was applied to pinpoint differential genes across two distinct disease states within a specific cell type (adjusted p-value < 0.05, and the absolute log2 fold change > 0.5). (**C**) The heatmap representing the gene expression of differentially expressed genes between T1D and HD in central memory CD4 TCM. (**D-E**) The GSEA results reveal TNF, NF-kb, and interferon signaling enrichment in CD4^+^ TCM in PLN (**D**) and spleen (**E**). (**F**) The plot indicating pre-ranked gene set enrichment analysis (GSEA) depicts the enrichment of TNF, NF-kb, and interferon signaling in CD4 TCM of PLN but not in spleen. (**G**) The heatmap showing the gene expression of the leading-edge genes for the NF-kb pathway extracted from GSEA output in PLNs. (**H**) The dot plot displaying the expression of NF-kb related genes and interferon genes combining cells across donors for each donor group. (**I**) The circle plots demonstrating outcomes of Immune Dictionary analysis, depicting the enrichment of top cytokines in CD4 TCM for upregulated differentially expressed genes obtained from T1D vs. HD and AAb^+^ vs. HD in PLNs, in addition to T1D vs. HD in spleen. Interferon cytokines, TNF, and NF-kb related cytokines are significantly enriched in PLNs of T1D and AAb^+^. T1D: Type 1 diabetes, AAb^+^: Autoantibody positive, HD: Healthy control donor. CD4 TCM: CD4^+^ central memory T cells. DEG: differentially expressed genes.

Next, we identified pathways enriched in differentially expressed genes, employing the gene set enrichment analysis (GSEA) strategy^26^. In PLN samples, the NF-kappaB (NF-kb) signal transduction in addition to type I and type II interferon pathways were selectively enriched among genes activated in central memory CD4^+^ T cells of T1D compared to HD donors (Figure 3D,F). The NF-kb signal transduction and type I interferon pathways were also enriched in central memory CD4^+^ T cells of AAb^+^ vs HD donors (Figure 3D). The interferon pathway has been implicated in many autoimmune diseases including T1D^27,28^. In spleens, B cells but not CD4^+^ T cells demonstrated enrichment of TNF signaling and NF-kb pathways in the comparison of T1D vs HD donors (Figure 3E). Visualization of genes at the leading-edge of the NF-kb gene-set used in the GSEA analysis further confirmed their differential expression across donors in PLNs (Figure 3G). Dot plot analysis of mRNA levels for genes that encode the NF-kb signaling component such as *RELA, NFKB1, RELB* in addition to the TNF-a receptors including *TNFRS1A* and *TNFRS1B* (also called *TNFR1* and *TNFR2*) in central memory CD4^+^ T cells clearly demonstrated their activation in T1D and AAb^+^ donors (Figure 3H). We performed a separate analysis for samples collected from the head of the pancreas and were able to recapitulate the upregulation of NF-kb associated pathways in T1D and AAb^+^ donors (Figure S3D-F). In sum, genes induced by TNF in addition to genes induced by interferons were significantly more expressed in central memory CD4^+^ T cells residing in PLNs of T1D and AAb^+^ donors.

Our GSEA suggested the increased expression of TNF and interferon target genes in PLNs of T1D and AAb^+^ donors. However, gene sets available in databases such as KEGG^29^ and REACTOME^30^ originate from broad biological contexts and fail to describe cell-type-specific cytokine effects. To infer cytokines interacting specifically with central memory CD4^+^ T cells in PLN tissues of T1D donors, we leveraged data from the “Immune Dictionary”, which is a resource with single-cell transcriptomic profiles of more than 17 immune cell types in response to 86 cytokines in mouse lymph nodes *in vivo*^31^. The Immune Response Enrichment Analysis (IREA) in this resource provides a link between cytokines and specific cell types based on a statistical test performed on experimental data. Although cytokine stimulation and transcriptional profiling of Immune Dictionary are produced in mice, the information gained is applicable to human cells^31^. The integration of our differentially expressed genes with the Immune Dictionary predicted that interferon alpha, beta, gamma, IL36α^32^ (an IL1 family cytokine that can activate NF-kb), IL18 (another IL1 family cytokine that can activate NF-kb), IL12, TL1A^33^ (a TNF family cytokine that can activate NF-kb), and TNF-a (the prototypic TNF family cytokine) are the most likely cytokines causing changes in gene expression in central memory CD4^+^ T cells of PLNs from T1D donors. Moreover, it was predicted that TL1A, IL36a, and TNF-a in central memory CD4^+^ T cells induced the expression of genes that were upregulated in PLN tissues of AAb^+^ vs HD donors (Figure 3I). A similar analysis in spleens suggested that a different combination of cytokines namely IL7, FLT3L, IL36a, IL-12, VEGF, and IL23 acting on central memory CD4^+^ T cells of T1D donors (Figure 3I). Altogether, our analysis suggests distinct cytokine environments in PLNs vs spleens of T1D donors, reporting a strong activation of TNF cytokine family members inducing the expression of the transcription factor NF-kb and its target program.

### Large-scale chromatin remodeling in central memory CD4^+^ T cells in T1D PLNs

The spatiotemporal control of gene expression is achieved through the combinatorial activity of cis-regulatory elements (CREs) such as enhancers and promoters distributed across the genome. DNA sequences encoding CREs undergo a complex process of epigenetic regulation such as DNA methylation, chromatin accessibility, and histone modifications, which license CREs to engage in gene regulation^34^. Mapping chromatin accessibility using ATAC-seq at single-cell and bulk levels has established itself as a reliable strategy to profile candidate CREs (cCREs)^34^. The salient advantage of our multiome assay is that the ATAC modality can be used to map cCREs while the RNA modality can reliably annotate cell types. We observed that from 108,715 chromatin accessibility peaks identified in our ATAC modality, 80% overlapped ENCODE cCREs^35^ and ∼10% overlapped promoters (Figure S4A-B), suggesting the high quality of our chromatin accessibility measurements.

To delineate cCREs with deregulated accessibility across T1D donors, we performed pseudo-bulk differential accessibility analysis between donor groups for each annotated cell type in PLNs and spleens, separately. Using stringent criteria (|log2 fold change|>1 and p-value<0.05), we reported the number of cCREs whose accessibility increased or decreased in the pair-wise comparisons between T1D vs HD, T1D vs AAb^+^, and AAb^+^ vs HD donors (Figure 4A-B). Recapitulating changes in gene expression, we found the largest number of differentially accessible cCREs between T1D and HD donors (Figure 4A-B). Although cells from AAb^+^ and HD donors also reflected differences in open chromatin regions, the T1D vs AAb^+^ comparison demonstrated minimal number of differentially accessible cCREs for any cell type, implying a high degree similarity in chromatin landscapes of T1D and AAb^+^ donors (Figure 4A-B). Remarkably, in agreement with our findings from the gene expression analysis, the comparison of various cell types in PLN samples highlighted central memory CD4^+^ T cells as the cell type exhibiting the most dramatic changes in chromatin accessibility between T1D and HD donors (Figure 4A). The heatmap visualization of chromatin accessibility levels at differential cCREs from central memory CD4^+^ T cells in T1D vs HD comparison also corroborated their differential accessibility by the donor state (Figure 4C). Although most cCREs in this group did not pass the statistical threshold to be referred as differentially accessible between AAb^+^ and HD donors, the heatmap visualization suggests a high degree of similarity between AAb^+^ and T1D donors (Figure 4C). In contrast to PLNs, memory B cells in spleens reflected the largest difference in chromatin accessibility levels in T1D vs HD comparison (Figure 4B).

**Figure 4.**
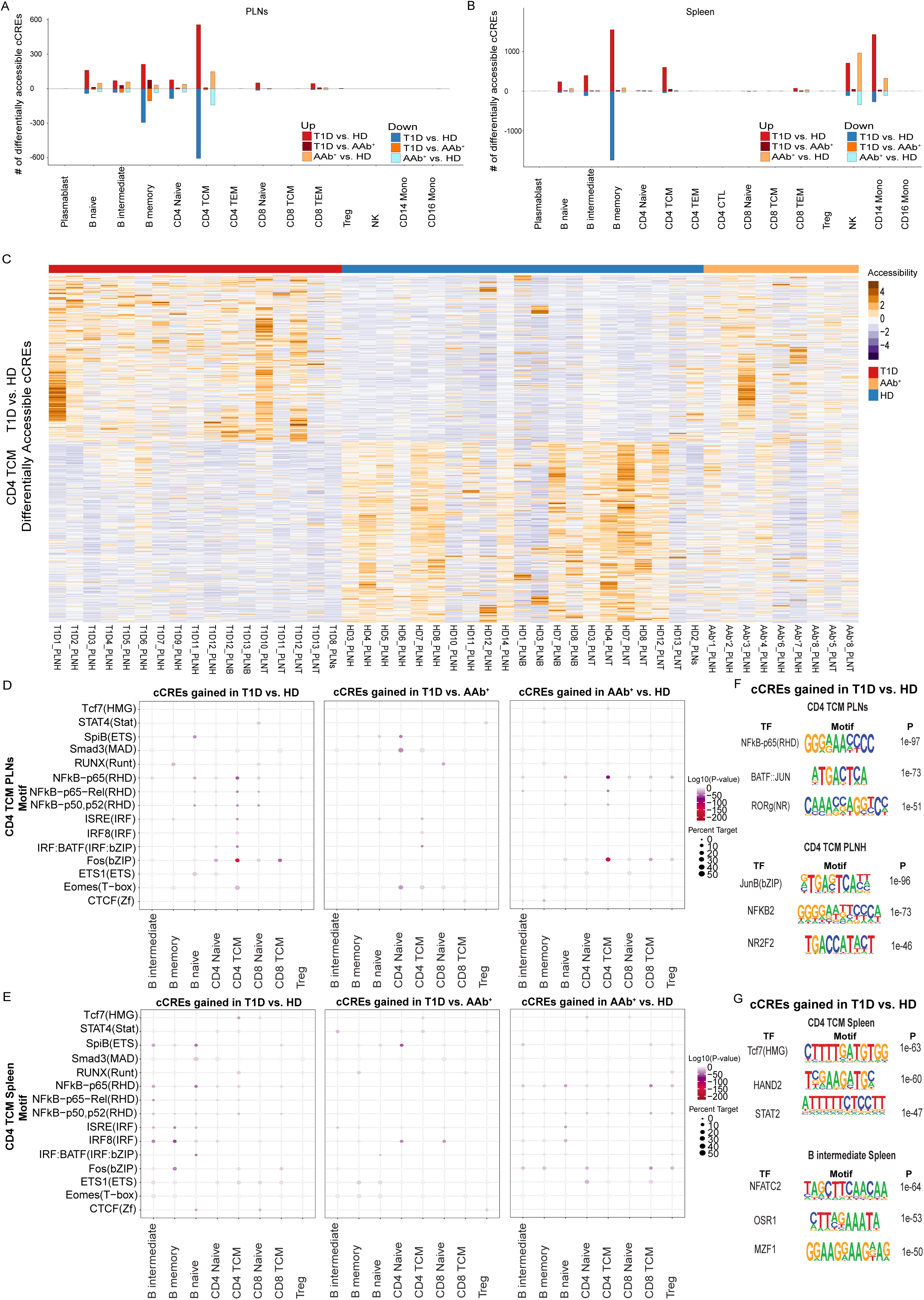
Central memory CD4^+^ T cells are the most epigenetically affected immune cells in T1D PLNs. (**A-B**) The bar plot demonstrating the number of differentially accessible cCREs in each cell type between T1D vs HD, T1D vs AAb^+,^ and AAb^+^ vs HD in PLNs (**A**) and spleen (**B**). (**C**) The heatmap representing chromatin accessibility levels for differentially accessible cCREs between T1D and HD in CD4 TCM. (**D-E**) Motif analysis reveals top transcription factors enriched for each cell type for PLN (**D**) and spleen (**E**). (**F-G**) Seqlogo depicting *de novo* motif analysis using Homer in gained accessible chromatin regions in CD4 TCM of T1D donors compared with HD in PLNs (**F**), and CD4 TCM and B intermediate cells of T1D donors compared with HD in spleen (**G**). T1D: Type 1 diabetes, AAb^+^: Autoantibody positive, HD: Healthy control donor. DA: Differential accessible, PLNH: pancreatic lymph node from the pancreas head. CD4 TCM: central memory CD4^+^ T cells.

We noted that the two non-diabetic donors, AAb^+^ #3 and #7 exhibited chromatin accessibility at these cCREs similar to T1D donors (Figure 4C). A closer look into differentially expressed genes across donors also confirmed similarity of gene expression for these two individuals and T1D donors (Figure 3C,G). Notably, high levels of glutamic acid decarboxylase (GAD) and Islet Antigen 2 (IA-2) autoantibodies were detected in donor AAb^+^ #7, and a high level of GAD autoantibody was detected in donor AAb^+^ #3 (Table S1), suggesting that these individuals might have been further along the path towards T1D than other non-diabetic AAb^+^ organ donors. In sum, differential accessibility analysis of cCREs using the ATAC modality recapitulated findings of the RNA modality and confirmed that central memory CD4^+^ T cells are the most dramatically changed cell type in PLNs of T1D and AAb^+^ donors compared to the HD group.

To identify transcription factors involved in remodeling the chromatin landscape of cells residing in PLNs of T1D and AAb^+^ donors, we performed motif analysis within differentially accessible cCREs for each cell type. Curating a list of known motifs related to transcription factors expressed in immune cells and searching the enrichment of their recognition sites within cCREs, we determined the strong enrichment of NF-kb-p65, NF-kb-p50, interferon-sensitive response element (ISRE) and AP-1 motifs in cCREs selectively accessible in central memory CD4^+^ T cells of T1D vs HD donors (Figure 4D). A similar enrichment for NF-kb-p65, NF-kb-p50, and AP-1 motifs was detected in cCREs selectively accessible in central memory CD4^+^ T cells of AAb^+^ vs HD donors, suggesting that NF-kb and interferon-associated transcription factors are critical in chromatin remodeling of T1D and AAb^+^ donors. We did not detect this enrichment in central memory CD4^+^ T cells of spleens. Instead, the naïve B cells present in the spleen of T1D and AAb^+^ donors exhibited increased accessibility at cCREs with enrichment of NF-kb-p65, IRF-BATF and SpiB (ETS) transcription factors (Figure 4E). *De novo* motif analysis and seq-logo visualization of cCREs gained in T1D compared with HD donors corroborates selective enrichment of NF-kb and AP-1 motifs in central memory CD4^+^ T cells residing in PLNs but not spleens (Figure 4F-G). We performed a separate analysis for PLN samples collected from the head of the pancreas and were able to recapitulate chromatin remodeling at cCREs enriched with binding sites for NF-kb associated genes in T1D and AAb^+^ donors (Figure S4C-G). Since the chromatin structure is an attractive template for both transmitting and remembering cytokine signaling^17,18^, our analysis presents *de novo* chromatin remodeling at different stages of T1D progression by NF-kb transcription factors, which mediate the nuclear response to TNF-a, IL36a, and TL1A.

### Association of T1D risk variants and chromatin accessibility of immune cells

Large-scale mapping of cCREs in peripheral blood of healthy donors by consortiums such as ENCODE and Epigenomics Roadmap projects clearly demonstrated the enrichment of SNPs associated with autoimmune diseases within cCREs of immune cells^36,37^. The interpretation of these findings has been that disruption of regulatory elements in immune cells of individuals with risk variants leads to changes in gene expression, causing autoimmunity. However, chromatin accessibility maps of affected tissues in individuals diagnosed with autoimmune disorders have rarely been available. The ATAC modality in our multiome measurements allowed us to assess the enrichment of T1D SNPs within cCREs which gained or lost accessibility in T1D donors in a cell type-specific manner. Our statistical analysis relying on a multiple regression model^38^ revealed the strongest enrichment of T1D SNPs occurs within cCREs of central memory CD4^+^ T cells whose accessibility increased in T1D donors compared with HD (Figure 5A). As expected from earlier studies reporting a large cluster of immune-mediated diseases forming a complex network of shared genetic loci^37^, we detected a similar enrichment of T1D-specific cCREs in SNPs associated with other immune diseases such as rheumatoid arthritis. We also found the highest enrichment of T1D SNPs within cCREs of Tregs whose accessibility decreased in T1D donors compared with HD, suggesting a novel link between genetics of T1D and loss of tolerance (Figure 5B). However, we did not observe any significant enrichment of risk loci for diseases with limited immune contribution such as T2D and Alzheimer’s disease in cCREs whose accessibility changed in T1D (Figures 5C-D and S5A-D). Together, T1D associated variants frequently fall within cCREs that selectively gained accessibility in central memory CD4^+^ T cells residing in T1D PLNs.

**Figure 5.**
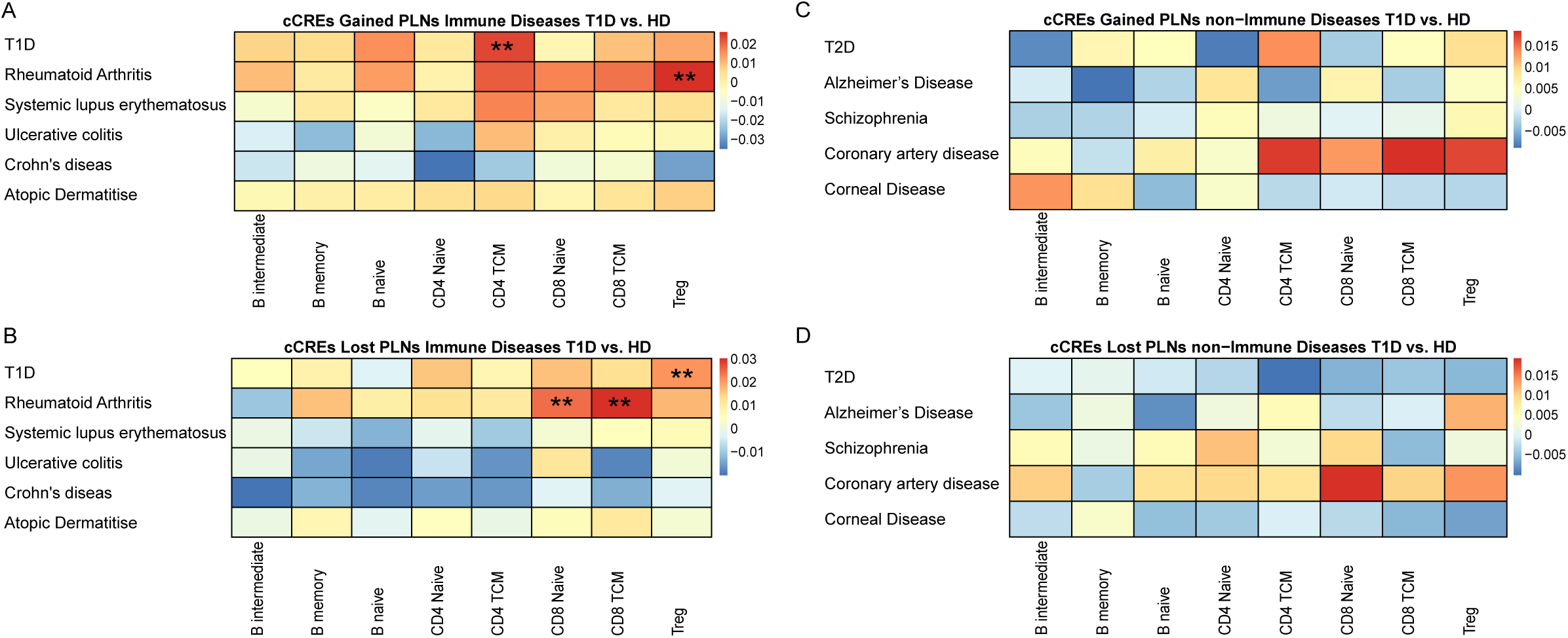
T1D-associated SNPs were significantly enriched in gained accessible regions of CD4^+^ central memory T cells. (**A-D**) The enrichment of T1D and other immune disease-associated SNPs in cCREs gained T1D vs. HD (**A**) and lost in T1D vs. HD (**B**). The enrichment of non-immune disease-associated SNPs in in cCREs gained T1D vs. HD (**C**) and lost in T1D vs. HD (**D**).

### Single-cell transcriptional profiling in NOD mice corroborates the enrichment of TNF pathway and NF-kb genes during T1D progression

Joint profiling of gene expression and chromatin accessibility in PLNs of human T1D organ donors revealed the aberrant expression of TNF target genes including the induction of NF-kb and the *de novo* chromatin remodeling of NF-kb associated cCREs in central memory CD4^+^ T cells. Next, we examined whether these findings can be validated in the non-obese diabetic (NOD) mouse strain, the mouse model of spontaneous autoimmune diabetes. In NOD animals, a marked decrease in pancreatic insulin content occurs in females at about 12 weeks of age and several weeks later in males. We isolated PLNs from female NOD mice at 2, 4, and 9 weeks of age, representing the pre-symptomatic stage. We also performed similar experiments in 20 weeks old females, selecting mice in which diabetes had been confirmed by increased blood and urine glucose levels (Figure 6A). To study the relevance of TNF family pathway at early T1D development in this mouse model, we performed single-cell RNA-seq experiments in CD4^+^ T cells purified from PLNs of these mice. Following a standard single-cell RNA-seq analysis^39^, we annotated clusters based on the expression of marker genes grouping CD4^+^ T cells in naïve, activated, regulatory (Treg), and proliferating states (Figures 6B-D and S6A-C). Although CD4^+^ Tregs from all ages formed a single cluster, naïve CD4^+^ T cells from 2 weeks old and activated CD4^+^ T cells from T1D mouse of 20 weeks old formed distinct clusters, reflecting their transcriptomic differences (Figure 6D). Strikingly, when we compared activated CD4^+^ T cells from 20 weeks old mice with naive CD4^+^ T cells from 2 weeks old mice via the Immune Response Enrichment Analysis introduced above, we found an enrichment for IL36a, interferon-alpha, IL12, TL1A, interferon-gamma, and IL18. In line with this analysis, NF-kb signature genes were also expressed at various stages of T1D progression in NOD mice including the pre-symptomatic stages (Figure 6E-G). Taken together, these data corroborate our findings in human PLNs of AAb^+^ and T1D organ donors.

**Figure 6.**
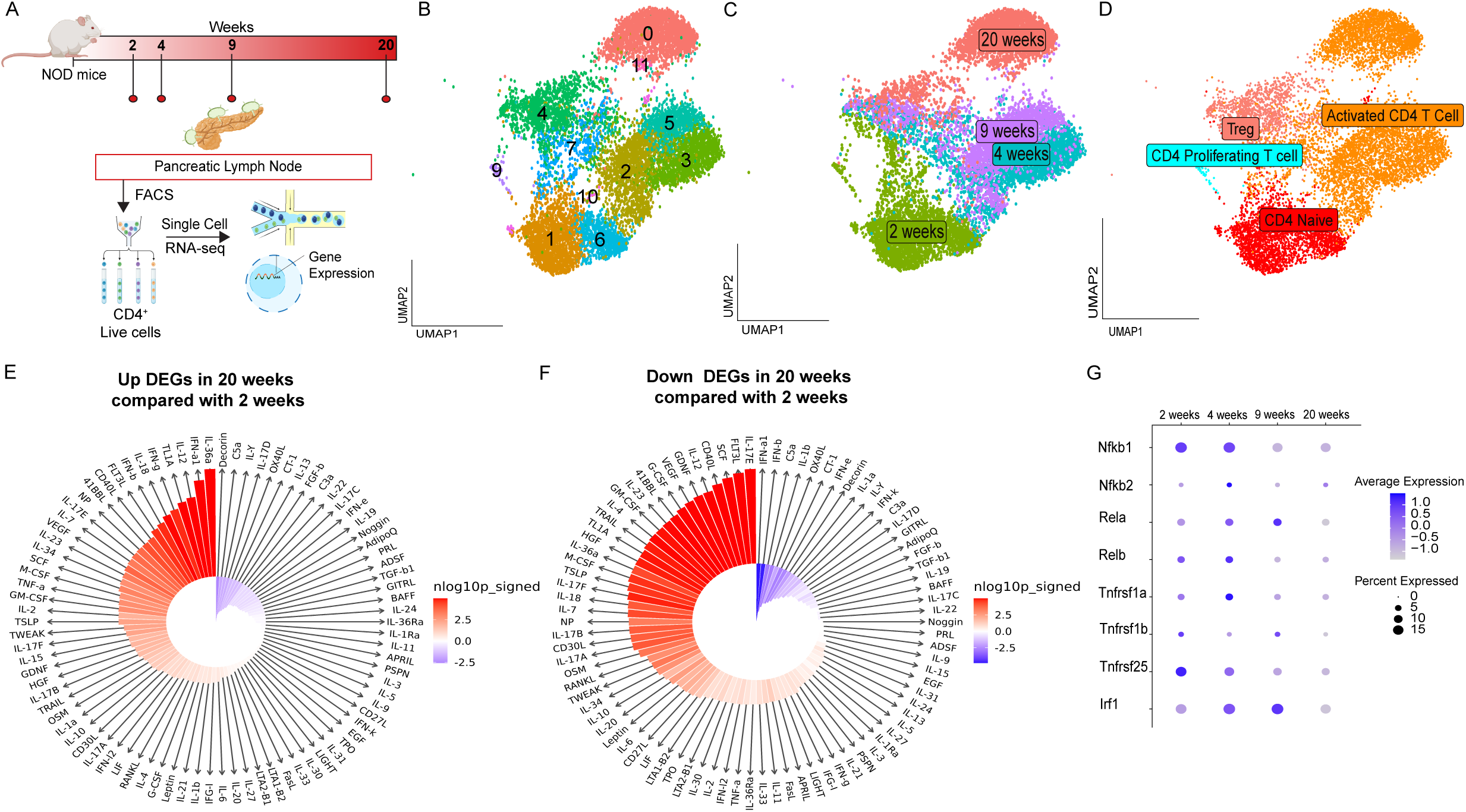
TNF and interferon-related genes are upregulated in the early stages of T1D in PLNs of NOD mice. (**A**) Schematic outlining the experimental approach for NOD mice. (**B-D**) The Uniform Manifold Approximation and Projection (UMAP) representing immune cells remained after quality control analysis labeled with cluster number (**B**), tissue time point (**C**), and immune cell annotation (**D**). The circle plots demonstrating the enrichment of top cytokines in CD4^+^ for upregulated DEGs (E-H). DEGs: Differentially expressed genes.

## Discussion

Cytokines encompass a diverse group of small proteins that are secreted and act mostly locally by binding to specific receptors on target cells^31^. This receptor-ligand interaction transmits cellular signals to the nucleus by activating transcription factors which can alter the chromatin landscape and turn on the gene programs required for coordinating various activities among different cell types within the immune system. It is not surprising that cytokine-based approaches have been in use for decades for the treatments of autoimmune diseases. Ongoing clinical trials for early onset or at risk T1D individuals are investigating immunotherapeutic approaches aimed at blocking inflammatory cytokines such as interferons and TNF. Yet, a substantial gap remains in our understanding of the precise timing and cellular mechanisms through which these cytokines contribute to T1D progression. One strategy to fill these gaps is to perform large-scale immune profiling in humans. However, deep immune profiling studies for human T1D have mostly been performed on leukocytes from the peripheral blood^10,27,40^, which is not the site of pathogenesis. Hence, blood-based immune profiling fails to uncover local cytokine events related to T1D progression. Our objective in this study was to determine the identity of immune subsets receiving aberrant cytokine signals during T1D progression using molecular profiling of immune cells in PLNs. The rationale for deep profiling of PLNs is that priming of autoreactive T cells occurs at these secondary lymphoid organs^15,22^. Supporting this notion, excision of PLNs in NOD mice almost completely protect young mice against diabetes development^14^. Our work represents the largest and most advanced molecular measurements of immune events in pancreatic tissues for human health and autoimmunity. Our fully annotated multiome data along with extensive clinical and metabolic information can be accessed through PANC-DB (https://hpap.pmacs.upenn.edu/), which is HPAP’s open-access database.

The central finding of our large-scale immune profiling is that central memory CD4^+^ T cells in PLNs but not spleens of T1D donors exhibit increased expression of the proinflammatory transcription factor NF-kb and chromatin remodeling associated with this transcription factor. Remarkably, this observation was also true for AAb^+^ donors even though 7 out of 8 AAb^+^ donors only express one antibody i.e. GAD, indicating the association of NF-kb and its downstream targets in this group. The *de novo* nucleosome remodeling mediated by NF-kb in CD4^+^ T cells was also linked with T1D SNPs, implying the role of T1D genetics in this chromatin signature. A major advantage of measuring gene expression and chromatin accessibility at the same time is the ability to assess the relevance of upstream factors mediating signal transduction to the nucleus. It has been recently shown that the transcription factor NF-kb orchestrates *de novo* nucleosome remodeling during the primary response to Toll-like receptor 4 signaling in macrophages^41^. Our integrative data analysis of gene expression and chromatin accessibility implies the role of cytokines inducing NF-kb in central memory CD4^+^ T cells including TNF-a, TL1A and IL36a.

To further validate our findings, we performed single-cell RNA profiling in the experimentally tractable NOD mouse model which enables the analysis of disease progression along a predictable time course. It is worth pointing out that TNF inhibition has been a major topic of investigation in this animal model for decades. In 1998, Flavell and colleagues reported that transgenic expression of TNF-a promotes diabetes in NOD mice^42^. This observation was followed by other groups including von Herrath and colleagues who reported a dual role for TNF-a in NOD mice^43^. They showed while blockade of TNF-a during an early phase of T1D in young NOD animals prevented diabetes completely, TNF-a expression enhanced the autoimmune process when induced late^43^. These data certainly imply the pro-inflammatory role of TNF cytokine at early stages of autoimmunity in agreement with the early expression of NF-kb genes in our single-cell profiling in mice. Our study remains the first comprehensive immune profiling in PLNs of human T1D, reporting selective induction of TNF target genes in central memory CD4^+^ T cells. Our data strongly suggest that golimumab, a TNF-a inhibitor currently undergoing phase II clinical trials to delay T1D onset in at-risk individuals, may operate by inhibiting central memory CD4^+^ T cells in PLNs, thereby mitigating the autoimmune interaction of CD8^+^ T cells with beta cells. Our study provides a mechanistic rationale for the efficacy of TNF-a inhibitors and may motivate development of inhibitors for other cytokines that activate NF-kb signaling such as TL1A and IL36a, which were predicted to be critical to CD4^+^ T cell biology in T1D by our analysis. In the future, parallel immune cell profiling of peripheral blood and PLNs of T1D and AAb^+^ donors might yield biomarkers for living individuals.

